# Pneumonia induced rise in glucagon promotes endothelial damage and vascular thrombogenicity

**DOI:** 10.1101/2024.05.03.592488

**Authors:** Pegah Ramezani Rad, Vanasa Nageswaran, Lisa Peters, Leander Reinshagen, Johann Roessler, Szandor Simmons, Erik Asmus, Corey Wittig, Markus C. Brack, Geraldine Nouailles, Emiel P. C. van der Vorst, Sanne L. Maas, Kristina Sonnenschein, Barbara Verhaar, Robert Szulcek, Martin Witzenrath, Ulf Landmesser, Wolfgang M. Kuebler, Arash Haghikia

**Affiliations:** Department of Cardiology, Angiology and Intensive Care Medicine, Deutsches Herzzentrum der Charité, Campus Benjamin Franklin, Berlin, Germany; German Center for Cardiovascular Research (DZHK), Partner Site Berlin, Berlin, Germany; Friede Springer-Cardiovascular Prevention Center at Charité, Charité-Universitätsmedizin Berlin, Berlin, Germany; Institute of Chemistry and Biochemistry, Freie Universität Berlin, Berlin, Germany; Department of Cardiology, University Hospital St Josef-Hospital Bochum, Ruhr University Bochum, Bochum, Germany; Institute of Biology, Freie Universität Berlin, Berlin, Germany; Institute of Physiology, Charité-Universitätsmedizin Berlin, corporate member of the Freie Universität Berlin and Humboldt-Universität zu Berlin, Berlin, Germany; Laboratory of in vitro modeling systems of pulmonary and thrombotic diseases, Institute of Physiology, Charité – Universitätsmedizin Berlin, corporate member of Freie Universität Berlin and Humboldt-Universität zu Berlin, Charitéplatz 1, 10117 Berlin, Germany; German Center for Lung Research (DZL), Partner site Berlin, Germany; Department of Infectious Diseases, Respiratory Medicine and Critical Care, Charité-Universitätsmedizin Berlin, corporate member of the Freie Universität Berlin and Humboldt-Universität zu Berlin, Berlin, Germany; Institute for Molecular Cardiovascular Research (IMCAR), RWTH Aachen University, 52074 Aachen, Germany; Aachen-Maastricht Institute for CardioRenal Disease (AMICARE), RWTH Aachen University, 52074 Aachen, Germany; Institute for Cardiovascular Prevention (IPEK), Ludwig-Maximilians-University Munich, 80336 Munich, Germany; Interdisciplinary Center for Clinical Research (IZKF), RWTH Aachen University, 52074 Aachen, Germany; Department of Cardiology and Angiology, Hannover Medical School, Hannover, Germany; Institute of Molecular and Translational Therapeutic Strategies (IMTTS), Hannover Medical School, Carl-Neuberg-Str. 1, 30625, Hannover, Germany; Department of Vascular Medicine, Amsterdam University Medical Centers (UMC), Amsterdam, The Netherlands; Department of Public and Occupational Health, Amsterdam University Medical Centers (UMC), Amsterdam, The Netherlands; The Keenan Research Centre for Biomedical Science at St. Michael’s, Toronto, Canada; Departments of Surgery and Physiology, University of Toronto, Toronto, Canada

## Abstract

**Background:** Recent studies have demonstrated a link between respiratory infections and increased short-term risk of cardiovascular disease (CVD). However, the molecular mechanisms underlying the increased cardiovascular risk after respiratory infections are only poorly understood. Here, we aimed to decipher pathophysiological circuits of pneumonia associated CVD in experimental models of bacterial pneumonia and vascular injury.

**Methods:** C57BL/6J mice were exposed to intranasal inoculation with either *Streptococcus pneumoniae* (*S. pneumoniae*) serotype 4 (pneumonia group) or phosphate buffered saline (PBS) (control group). 24 hours *post infectionem* (p.i.) mice were treated with antibiotics until the end of the study. On day 7 p.i. carotid artery injury (CI) was induced by electric stimulation and vascular repair was analyzed 3 days after injury. Plasma proteomic analyses were performed by Olink Bioscience. Primary human aortic endothelial cells (HAECs) were used to study alterations of the endothelial functional properties, bioenergetic state and thrombogenic potential *in vitro*. Intravital fluorescence microscopy equipped with video recording was applied to measure thrombus formation in real-time.

**Results:** Bacterial pneumonia impaired repair capacity of the endothelium after vascular injury. Proteomic analyses revealed significantly higher plasma levels of glucagon in mice after recovery from pneumonia relative to controls, which was further confirmed by ELISA detecting glucagon. Mechanistically, we found that glucagon impaired mitochondrial bioenergetics and migratory potential in HAECs and induced an inflammatory response. Moreover, glucagon fostered vascular thrombogenicity as demonstrated by increased thrombocyte adhesion to HAECs and accelerated carotid artery thrombus formation *in vivo*. Acute application of the glucagon-like peptide-1 receptor (GLP1-R) agonist liraglutide to lower blood glucagon levels, restored vascular repair potential and attenuated vascular thrombogenicity in mice with pneumonia.

**Conclusions:** Our findings reveal a novel mechanism that associates elevated circulatory glucagon levels to dysfunctional endothelium and increased vascular thrombogenicity, suggesting glucagon signaling as a potential therapeutic target to prevent pneumonia-induced cardiovascular events.

**Graphical Abstract:** 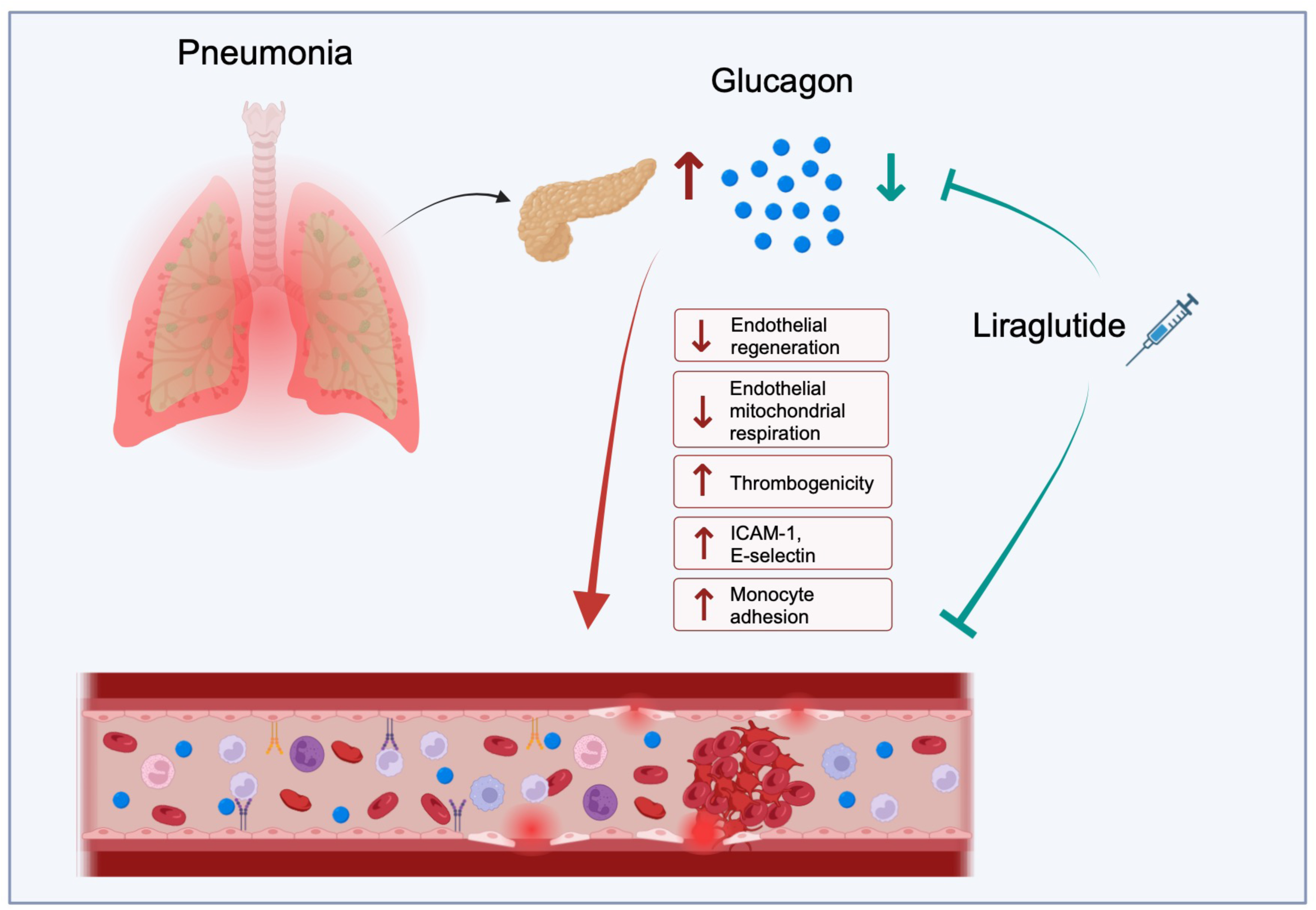

## Introduction

Community acquired pneumonia (CAP) is one of the most common medical diagnoses leading to hospitalization worldwide.^1^ Accumulating evidence from epidemiological studies points to a 2-fold to 8-fold increase in the risk of cardiovascular events, such as myocardial infarction and ischemic stroke, within the first 30 days after respiratory infections.^2,3^ For example, a Scottish observational study showed substantially increased incidence of myocardial infarction in the week following infection with *S. pneumoniae* or influenza virus, with age- and season-adjusted incidence ratios of 5.98 and 9.80, respectively. Conversely, vaccination against these infections has been shown to lower rates of myocardial infarction, supporting a causal link.^4^ Moreover, a causal link between CAP and increased risk of myocardial infarction is supported by studies reporting a significant decrease in the incidence of myocardial infarction upon vaccination against both *S. pneumoniae* infection^5^ and influenza virus infection.^6^

Although CAP has been an established acute trigger for cardiovascular events, the mechanistic links between CAP and increased risk of cardiovascular events still remains only poorly understood. However, detailed knowledge about CAP related factors contributing to cardiovascular events is essential for the development of preventive strategies, which may aid in reducing cardiovascular complications. Studies with experimental models of respiratory infections in atherosclerosis prone mice, as well as human autopsy studies of patients with systemic infections, have suggested activation of atherosclerotic lesions, resulting in increased vulnerability with elevated risk of coronary and cerebrovascular events^7,8^, although the detailed pathophysiological mechanisms underlying increased vascular vulnerability are still missing. In this study we focused on pathological alterations of the endothelium in an experimental model of pneumococcal pneumonia. In particular, we examined the effects of bacterial pneumonia on endothelial inflammation, thrombogenicity and regeneration after injury. The results of our study shed further insight into mechanistic circuits linking respiratory infections with cardiovascular events.

## Methods

### Data availability

The data, analytic methods, and study materials are available to other researchers for purposes of reproducing these results or replicating these procedures by reasonable request directed to the corresponding author.

### Mouse experiments

Wild type C57BL/6J mice were purchased from Janvier Labs (France). All mice experiments were performed at Campus Mitte Charité Cross Over (CCO) and were approved by the Ethics Committee on Animal Care and Use in Berlin, Germany. Mice were maintained under 12 hours light cycle in specific-pathogen-free conditions and all mice had access to food and water *ad libitum*.

To investigate the effect of bacterial pneumonia on arterial injury, male adult (17 weeks of age) C57BL/6J mice were randomly assigned to either control group, pneumonia group or liraglutide + pneumonia group. In pneumonia groups, mice were intranasally inoculated with 5 × 10^5^ *S. pneumoniae* TIGR4 in 20 µL PBS. Sham infection with 20 µL PBS was performed on control mice. Antibiotic treatment with ampicillin (0.4 mg/mouse) followed 24 hours p.i. every 12 hours by s.c. injections until day 5 p.i. Treatment was continued orally via drinking water until the end of the study with amoxicillin-clavulanic acid (6 mg/d/mouse). A third group was infected with *S. pneumoniae* as well and additionally treated with liraglutide (100 μg/kg BW), a GLP-1R agonist 0, 24 and 48 hours p.i. On day 7 p.i. carotid artery injury was performed in mice as described before^9^. To this end, mice were first anesthetized with 3% isoflurane and then anesthesia was maintained with 1% isoflurane in oxygen (1 L/min) via inhalation. Body temperature was maintained at 37°C using a heating pad to prevent hypothermia. The left side of the neck was shaved and a small incision was made, exposing the left carotid artery (LCA). Carotid injury was induced to the LCA by an electric impulse of 2 W for 2 seconds over a length of 4 mm using bipolar microforceps (VIO 50 C, Erbe Elektromedizin GmbH). Three days after carotid injury, endothelial regeneration was assessed by injecting 50 µL of 5% Evans blue (Sigma, cat no. E2129) into the left heart ventricle three minutes before sacrifice. The LCA was isolated, rinsed in PBS and fixed on microscope slides. The Evans blue-stained denuded area was captured *en face* by brightfield microscopy (Axioskop 40, Zeiss), and re-endothelialization was calculated as the ratio of white area to injured area using imaging software (ImageJ, NIH).

### Protein analysis Proteomics – Olink

Murine EDTA plasma was collected 10 days p.i. and *Target 96 Mouse Exploratory* proteomic panel, was performed by Olink Bioscience (Uppsala, Sweden). According to the manufacturer’s instructions using 1 µL plasma, protein profile was analyzed (n=5 per group). Sample quality control as well as normalization was performed using inter-plate controls. Normalized protein expression (NPX) values, Olink Proteomics’ arbitrary unit on log2 scale, was provided by Olink.

### Glucagon ELISA

For glucagon (Gcg) quantification murine whole blood samples were centrifuged at 800 x g for 15 min and plasma supernatant was transferred into a new 1.5 mL microfuge tube. 50 µL of murine plasma was then used to quantify glucagon concentrations with ELISA by following the manufacturer’s protocol (R&D Systems, cat no. DGCG0). Final glucagon concentrations were calculated based on a standard curve.

### Cell culture experiments

Cell culture experiments were carried out using primary human aortic endothelial cells (HAECs) (Cell Applications, cat. no 304-05a) in passage 8. HAECs were cultured in Endothelial Cell Growth Medium 2 (PromoCell GmbH, cat. No C-39211) which was supplemented with heat-inactivated 10% fetal calf serum (FCS) (c.c. pro, cat no. S-10-L), 100 units/mL penicillin and 100 µg/mL streptomycin (Gibco, cat no. 15140122). Human THP-1 monocytes were cultured in RPMI 1640 medium (Gibco, cat no. 11875093) supplemented with 10% FCS, 2 mM L-glutamine, 100 U/mL penicillin and 100 μg/mL streptomycin. Prior to experiments cells were incubated at 37°C in a 5% CO2 atmosphere and medium was changed every 2-3 days until cells were grown to confluence.

### Wound healing assay

*In vitro* scratch assay was performed to quantify the endothelial wound healing potential across a scratch-induced gap. HAECs (6 × 10^4^/well) were seeded on a fibronectin-coated 24-well culture plate overnight, and serum-starved (0.5% FCS) for 5 hours at 37°C in a cell culture incubator. An artificial gap was generated on the confluent cell monolayer using a sterile 200 µL pipette tip and cells were treated with 10 nM Gcg (Bachem, cat no. 4033017), 100 nM Gcg, or control for 16 hours. Pictures of the wound area were taken before stimulation and 16 hours thereafter on a phase-contrast microscope (EVOS XL Core; Thermo Fisher Scientific). The width of the scratch was measured and quantified by NIH ImageJ software.

### Flow cytometry of HAECs

The expression of the cellular adhesion molecules ICAM-1 and E-selectin was evaluated by flow cytometry. HAECs (6 × 10^4^/well) were seeded on 24-well plates and stimulated with 10 nM Gcg, 100 nM Gcg or control for 6 hours. Cells were washed, collected into FACS tubes (Falcon, cat no. 352052) and incubated with the following antibodies: AF700-labeled CD54 (1:100, BioLegend, cat no. 353126) and PE/Cy7-labeled CD62E (1:100, BioLegend, cat no. 336016) for 30 min at RT. Fluorescence-minus controls (FMOs) and unstained HAECs were included in the measurements. Samples were acquired using the Attune Nxt Flow Cytometer (Thermo Fisher Scientific) and analysis was performed with Kaluza software (Beckman & Coulter).

### Monocyte adhesion assay under flow

Adhesion of monocytes to endothelial cells was investigated by a flow-based adhesion assay. HAECs (2 × 10^5^ cells/slide) were seeded in ibidi y-shaped chamber slides (ibidi GmbH, cat no. 80126) for 5 hours and then incubated overnight under sterile flow conditions at a shear stress of 20 dyn/cm^2^. Cells were subsequently treated with 10 nM Gcg, 100 nM Gcg or control for 6 hours. Finally, Dil-labeled (Invitrogen, cat no. V22888) THP-1 monocytes (1 × 10^6^ cells/mL) were perfused through the chambers for 30 min. After incubation, non-adherent monocytes were removed with PBS and co-cultures were fixed with 4% paraformaldehyde (PFA). The number of adherent monocytes to HAECs was quantified from 12 different fields using a fluorescence phase-contrast microscope (BZ-X; Keyence Corporation).

### Endothelial blood flow assay

To assess platelet adhesion to endothelial cells, HAECs (24 × 10^3^/slide) were seeded at confluency in ibidi multichannel flow chamber µ-slides VI 0.4 (ibidi GmbH, cat no. 80606) and incubated at 37°C in a 5% CO2 atmosphere. After four days of incubation, confluent cells were stimulated with 10 nM Gcg, 100 nM Gcg or control for 6 hours. Whole blood stained with APC-tagged CD42b (1:500, Miltenyi Biotec, cat no. 130-100-208) was perfused over the cell monolayer in recalcification buffer (Tyrode’s buffer supplemented with 1% BSA and 2.1 mM Ca^2+^) for 5 min at a speed of 20 mL/h in accordance with our published protocol.^10^ Fifty µM adenosine diphosphate (ADP) (Sigma-Aldrich, cat no. A2754) was added directly to the perfusing of one channel per slide as positive control. At the end of the run slides were disconnected fixed with 4% PFA for 20 min at RT and washed with PBS. Before imaging, nuclei were stained with Hoechst 33342 (15 µg/mL; Miltenyi Biotec, cat no. 130-111-569) for 10 min at RT. The average percentage of CD42b covered area was quantified from five different fields per channel on a fluorescence phase-contrast microscope (EVOS M5000; Invitrogen).

### Seahorse assay

Mitochondrial function of HAECs was examined using a Seahorse XFe96 extracellular Flux analyzer (Agilent Technologies) by measuring oxygen consumption rates (OCR). In each well of XFe96-well microplates (Agilent Technologies, cat no. 102416-100), 15 × 10^3^ HAECs were plated and incubated with 10 nM Gcg, 100 nM Gcg or control for 6 hours. Cells were then washed with assay medium consisting of Seahorse XF DMEM Medium (Agilent Technologies, cat no. 103575-100) supplemented with 10 mM glucose (Carl Roth, cat no. X997.1) and 2 mM glutamine (Sigma-Aldrich, cat no. G3126). Next, 175 µL assay medium was added to each well and cells were incubated for one hour at 37°C in a CO2-free incubator. OCR was measured under basal conditions and in response to injections with 1 µM oligomycin (Sigma, cat no. 75351), 1 µM carbonylcyanid-4-(trifluormethoxy)phenylhydrazon (FCCP) (Sigma, cat no. C2920), 0.5 µM rotenone (Sigma, cat no. R8875) and 0.5 µM antimycin (Sigma, cat no. A8674). Wave software (version 2.6.0, Agilent Technologies) was used to calculate mitochondrial activity. Data were normalized for 1 × 10^3^ cells/well. For staining and normalization the assay medium was aspirated immediately after each assay and cells were washed with PBS, fixed with 4% PFA for 15 min and stained with 4′,6-diamidino-2-phenylindole (DAPI) (0.1 µg/mL; Roche, cat no. 10236276001) for 10 min at RT. A picture of each well was taken on a fluorescence phase-contrast microscope (Biorevo; BZ-9000; Keyence) and cells per well were counted using ImageJ Software (ImageJ, NIH).

### Lung histology

For histologic assessment, mice were prepared 48 hours and 10 days p.i. and lungs were carefully removed after ligation of the trachea to prevent alveolar collapse and fixed in 4% formaldehyde solution. Lungs were paraffin-embedded, cut into sections of 2 µm thickness and stained with hematoxylin and eosin (HE). To assess pathologic alterations, twenty pictures of evenly distributed horizontal sections of the entire lung were taken and evaluated according to a lung injury scoring system as previously described.^11^

### Thrombus formation assay using intravital microscopy

Mice were anesthetized by intraperitoneal (i.p.) injection of ketamine (200 mg/kg BW), xylazine (20 mg/kg BW) and NaCl in a total volume of 50-70 µL. Adequate depth of anesthesia was maintained throughout the whole experiment by continuous infusion of ketamine (200 mg/kg BW/hour) and xylazine (10 mg/kg BW/hour) and was controlled by regularly checking the intertoe reflex. The infusion rate of 10 mL/kg/h guaranteed fluid replacement throughout the imaging process. After anesthesia induction, 100 µL 0.1 mM rhodamine 6G (Carl Roth, cat no. 989-38-8) was injected via the tail vein, to fluorescently label platelets *in vivo*. To avoid hypothermia, anaesthetized mice were placed on a heating pad. The carotid arteries were surgically exposed and surrounding adipose tissue removed. In order to guarantee a local application of FeCl3 (Carl Roth, cat no. 7705-08-0) and to enable an even horizontal imaging axis, a black plastic tub was placed under the carotid artery. Imaging was performed on an intravital microscope (Transillumination, AxiotechVario, Carl Zeiss) equipped with a fibre-optic unit (Streppel Halolux 600), a CMOS camera (Sony ICE 6000, 1920 × 1080 pixels) and a BP 640/30 filter. After adding 2 µL 12.5% FeCl3 for 1 min using a filter paper, the remaining fluid was removed. Thrombus formation was then imaged using a 10x wet lens (Achroplan 10x/0.30 W Ph1, Carl Zeiss) and recorded at a frame rate of 50 frames per second (fps) for the selected vessel area of interest.

### Statistics

Statistical analyses for the *in vitro* and *in vivo* analyses were performed by GraphPad Prism 9 (GraphPad Software, Inc) and data are presented as mean ± standard error of mean (SEM). Gaussian distribution of data was assessed using the Shapiro-Wilk test and normally distributed data were analyzed by Student’s unpaired two-tailed t-test (for two groups) or one-way ANOVA followed by Bonferroni post hoc test (for ≥3 groups). For data significantly deviating from Gaussian distribution, nonparametric statistical analyses were performed using the Mann-Whitney U test for two groups or the Kruskal-Wallis test followed by Dunńs post hoc test for multiple comparisons (for ≥3 groups).

Proteomics data were analyzed in RStudio (v. 2023.9.1.494) using R (v. 4.2.1). Group differences were tested with Student’s unpaired two-tailed t-test. The significance level was set at p < 0.05. The R code was made publicly available in a Github repository (https://github.com/barbarahelena/olink-analyses-charite).

## Results

### Pneumococcal pneumonia impairs vascular regenerative potential after injury

A murine carotid injury model was used to investigate the effect of bacterial pneumonia on vascular regeneration after injury. As demonstrated in Figure 1A adult C57BL/6J mice were either infected with *S. pneumoniae* or PBS (control) and treated with antibiotics 24 hours p.i. until the end of the study. Lung histology confirmed lung injury in the acute phase of pneumonia 48 hours p.i. (Figure 1B) upon infection with *S. pneumoniae*. Notably, on day 10 p.i. lungs of mice infected with *S. pneumoniae* were not significantly injured, indicating effective treatment of bacterial pneumonia by antibiotic treatment (Figure 1B). Carotid artery injury was performed 7 days p.i. Quantification of Evans blue-stained area of the left carotid artery (LCA) determined the degree of de-endothelialization 3 days post injury (10 days p.i.). Our findings revealed that *S. pneumoniae* infection significantly impairs wound healing compared to control animals (Figure 1C).

**Figure 1.**
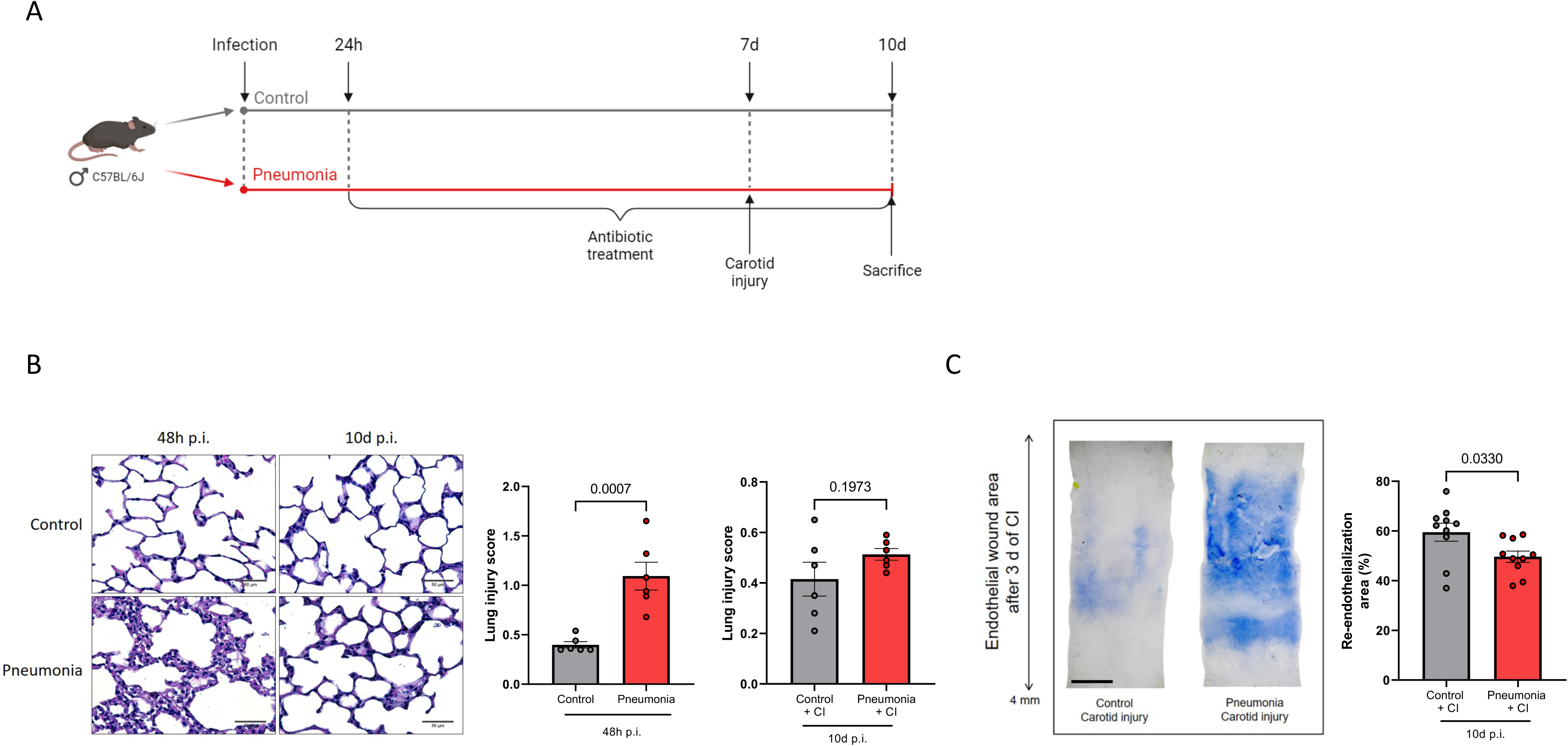
Bacterial pneumonia impairs endothelial regeneration in a murine carotid injury model. **A,** Experimental setup of C57BL/6J mice infected with *S. pneumoniae* and treated with antibiotics 24h p.i., followed by carotid artery injury (CI) to determine endothelial regeneration. **C**, Representative images of HE stained lungs from control mice and infected mice 48h and 10d p.i and quantification of lung injury score per timepoint. n=6 mice per group. 40x magnification, scale bar 50 µm. **D**, Representative images of Evans blue-stained murine carotid arteries from control and pneumonia groups three days after CI. The blue-stained area corresponds to injured area. The percentage of re-endothelialization was quantified as ratio between white-stained area/carotid injury area. n=10 mice per group. 5 x magnification, scale bar 500 μm.

### Murine plasma glucagon levels are increased upon pneumonia

In order to identify circulatory factors which may trigger the observed detrimental effects of bacterial pneumonia on vascular regeneration, the murine plasma proteome was profiled using the Olink proteomics platform. This analysis revealed a significant increase in glucagon levels among mice with bacterial infection (Figure 2A-C). ELISA analysis confirmed this finding showing a significant increase in circulating glucagon levels 10 days p.i (Figure 2D). Notably, plasma glucagon levels were not increased during the acute phase of pneumonia (48 hours p.i) (Figure 2E).

**Figure 2.**
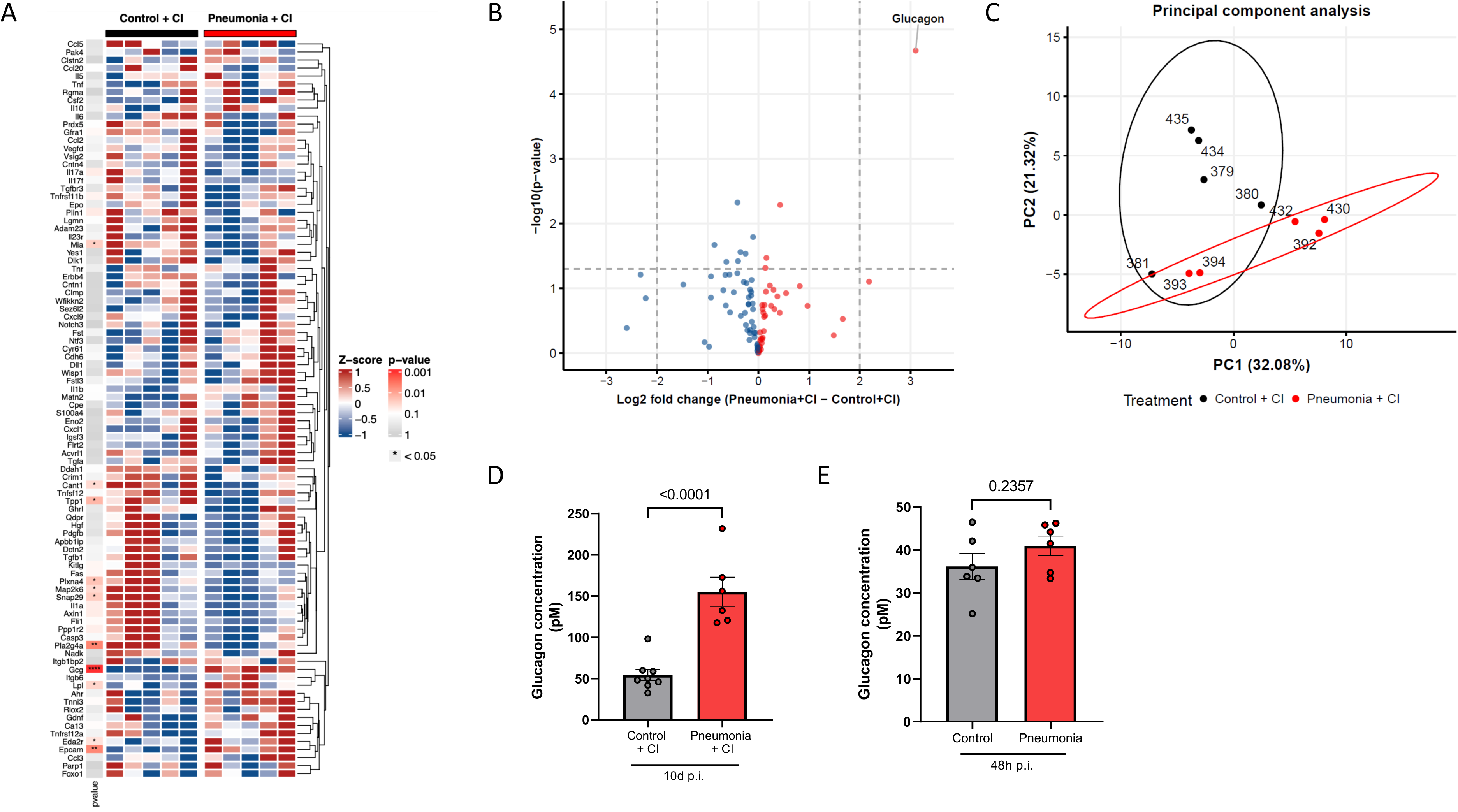
Pneumonia induces elevated circulatory glucagon concentrations. **A,** Heatmap of different levels of proteins measured by Olink. The heatmap color represents the normalized protein concentration (NPX). Red represents upregulated and blue represents downregulated protein levels. n=5 mice per group. **B**, Volcano plot demonstrates the log2 fold change (x-axis) versus -log10 p-value (y-axis) with the dashed line marking p = 1 × 10^−5^ for visual support. Each dot represents one of 92 proteins identified in murine EDTA plasma. n=5 per group. **C**, Principal component analysis of proteome profiles of each mouse. Black dots represents control group and red dots represent pneumonia group. n=5 per group. **D**, **E**, Glucagon ELISA shows glucagon concentration measured in pM 10d and 48h p.i. n=8 control mice 10d p.i and n=6 pneumonia mice 10d p.i. n=6 mice per each 48h p.i group.

### Glucagon impairs endothelial migratory capacity

To investigate a potential causal link between elevated glucagon levels after pneumonia and impaired vascular regenerative capacity, we examined the effect of glucagon on endothelial functional properties. To this end, we performed an *in vitro* wound healing assay using cultured HAECs. Our findings demonstrate that glucagon significantly impairs gap closure in the scratch-wound assay, suggesting compromised migratory capacity of glucagon-treated HAECs (Figure 3A and 3B). This observation indicates detrimental effects of glucagon on fundamental endothelial cell (EC) functions, that are crucial for the healing processes in the vascular wall after injury.

**Figure 3.**
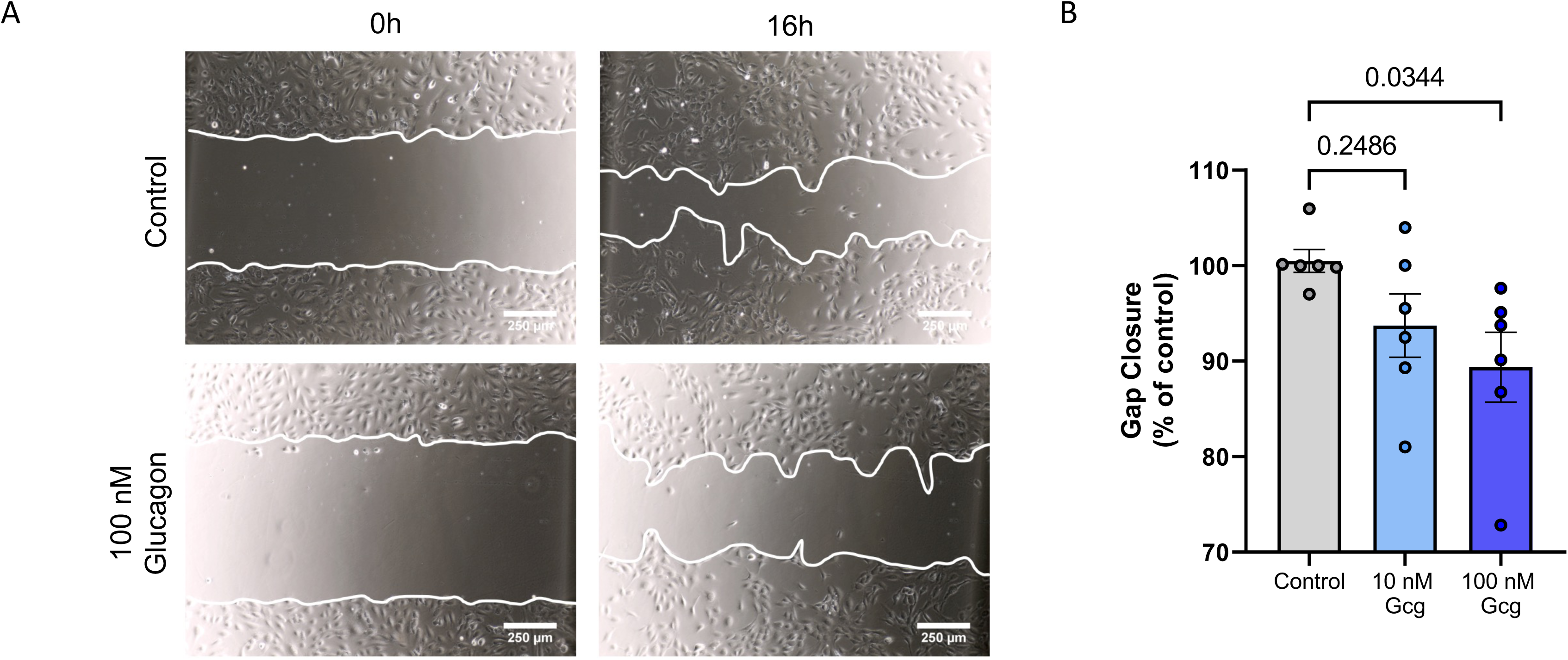
Effect of glucagon on endothelial migratory potential. **A**, Scratch-wound healing assay. Representative migration images of HAECs treated with 100 nM glucagon or control for 16 hrs. n=6 per group. 10 x magnification, scale bar 250 µm. **B**, Quantification of migration capacity. Gap closure was quantified as the width of the gap before versus after treatment.

### Glucagon impairs mitochondrial function of endothelial cells

In ECs the primary energy-producing mechanism is glycolysis, which takes place in the cytosol^12^. Complex mitochondrial networks in ECs^13^, however regulate the intracellular dynamics of NO, ROS and Ca^2+^, which in turn modulate endothelial cell function^14^. Thus, we assessed mitochondrial function in HAECs in response to glucagon stimulation to evaluate glucagon-mediated alterations of mitochondrial respiration. Endothelial OCR was therefore measured in real-time by Seahorse XFe96 extracellular Flux analyzer. Our results show significantly reduced maximal respiration and spare capacity upon glucagon stimulation (Figure 4B and 4C). The Seahorse Mito Stress Test mimics a physiological rise in energy demand in the mitochondria of HAECs by adding the uncoupler FCCP (Figure 4E). FCCP forces the respiratory chain to operate at maximum capacity through uninhibited electron flow along the electron transport chain. Cells stimulated with glucagon were not able to meet this metabolic challenge and had significantly reduced maximal respiration as compared to control cells, indicating impaired mitochondrial function (Figure 4B). This finding was accompanied by significantly reduced spare capacity upon glucagon stimulation, indicating reduced capability to respond to an energetic demand and thus being more sensitive to oxidative stress (Figure 4C). Basal respiration and ATP production did not change in response to glucagon (Figure 4A and 4D). In sum, these findings suggest glucagon-mediated impairment of mitochondrial function in metabolically challenged ECs, yet not at rest.

**Figure 4.**
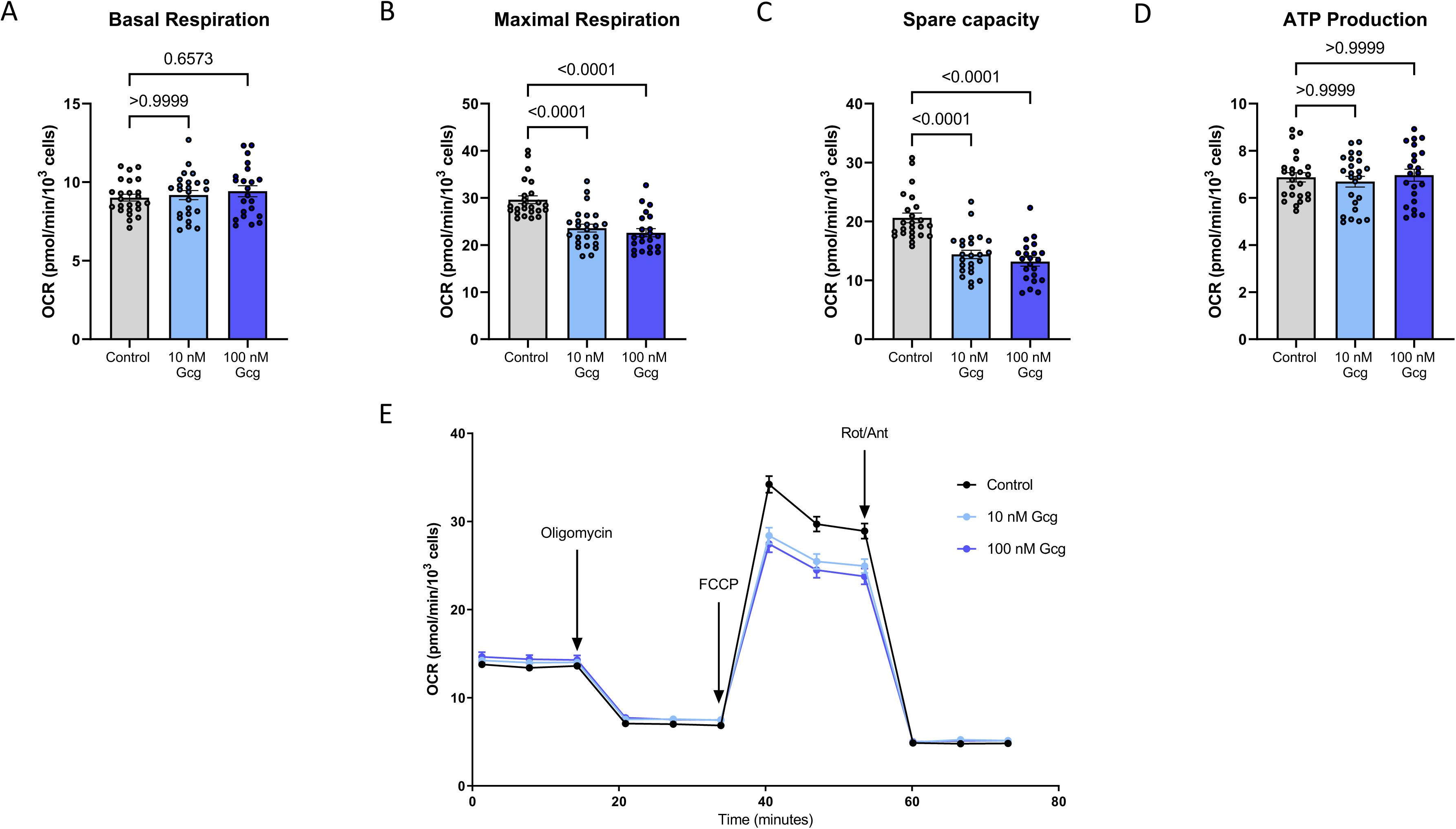
Mitochondrial respiration is impaired by glucagon. Measurement of real-time OCR in Seahorse mito stress test was performed (**E**) and basal respiration (**A**), maximal respiration (**B**), spare capacity (**C**) and ATP production (**D**) were calculated 6 hrs after glucagon stimulation. OCR measurements were normalized to 10^3^ cells per well. n=3 biological replicates.

### Glucagon promotes thrombogenicity

Since pneumonia is associated with increased atherothrombotic risk, we sought to determine the effect of glucagon on endothelial thrombogenicity. To this end, HAECs were pre-stimulated with glucagon for 6 hours and subjected to whole blood flow for 5 minutes, which showed significantly higher platelet adhesion as compared to vehicle treatment, demonstrating increased platelet-endothelial interactions following glucagon stimulation (Figure 5A and 5B). Stimulation with ADP served as positive control. To examine whether mice with pneumonia and elevated plasma glucagon levels were similarly at increased risk for thrombus formation, intravital microscopy was performed to measure the time to complete thrombotic vessel closure at the injured side of the carotid artery. In mice with pneumonia, thrombus formation and vessel closure occurred significantly faster as compared to control animals (Figure 6A-C). Next, we treated mice with the glucagon-like peptide-1 receptor agonist liraglutide to lower blood glucagon levels post-infection. Interestingly, thrombus formation time was restored upon treatment with liraglutide (Figure 6A-C). To evaluate whether glucagon in addition to increasing vascular thrombogenicity, also impacts platelet aggregation, whole blood samples from healthy donors were stimulated with glucagon and thrombus formation was quantified by aggregometry (Figure S1A). However, glucagon stimulation did not alter thrombocyte aggregation in whole blood. Moreover, the expression of platelet activation markers (glycoprotein IIb/IIIa, P-selectin and CD63) in whole blood samples were not altered upon stimulation with glucagon as compared to controls (Figure S2A-D). These findings demonstrate that pneumonia-associated elevation of blood glucagon levels impacts thrombogenicity on endothelial cells and promotes thrombus formation in the absence of direct effect on platelets.

**Figure 5.**
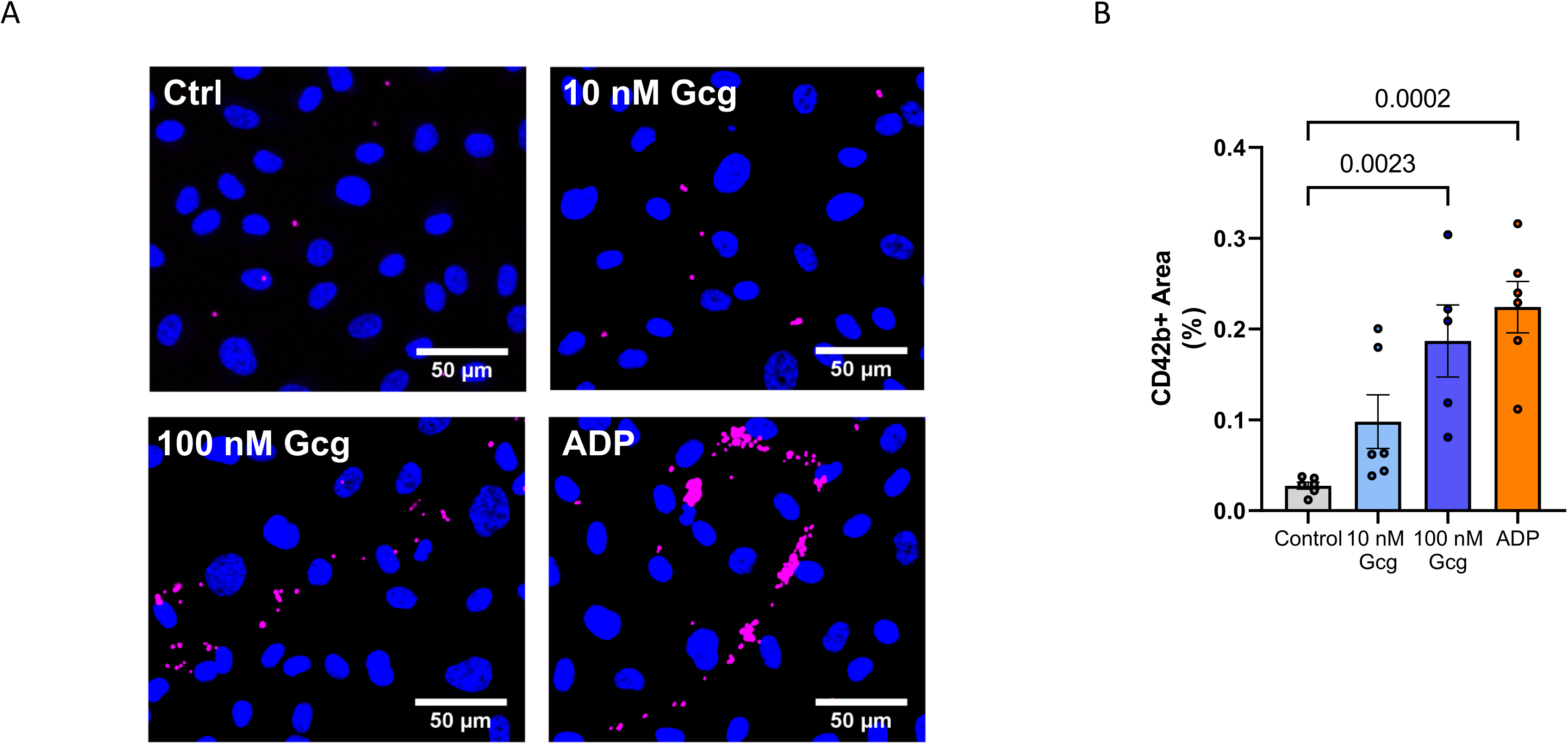
Glucagon leads to increased platelet adhesion to HAEC monolayers after perfusion with whole blood *in vitro*. **A**, Representative images of CD42b^+^ platelets (pink dots) adhered to HAECs after 6 hrs of glucagon stimulation, followed by 5 min of whole blood flow at 20 mL/h. Endothelial nuclei were counterstained with Hoechst 33342 (blue). Scale bar 50 µm. **B**, Quantification of CD42b^+^ coverage on HAEC monolayer. n=5 100 nM glucagon, n=6 control, 10 nM glucagon and ADP.

**Figure 6.**
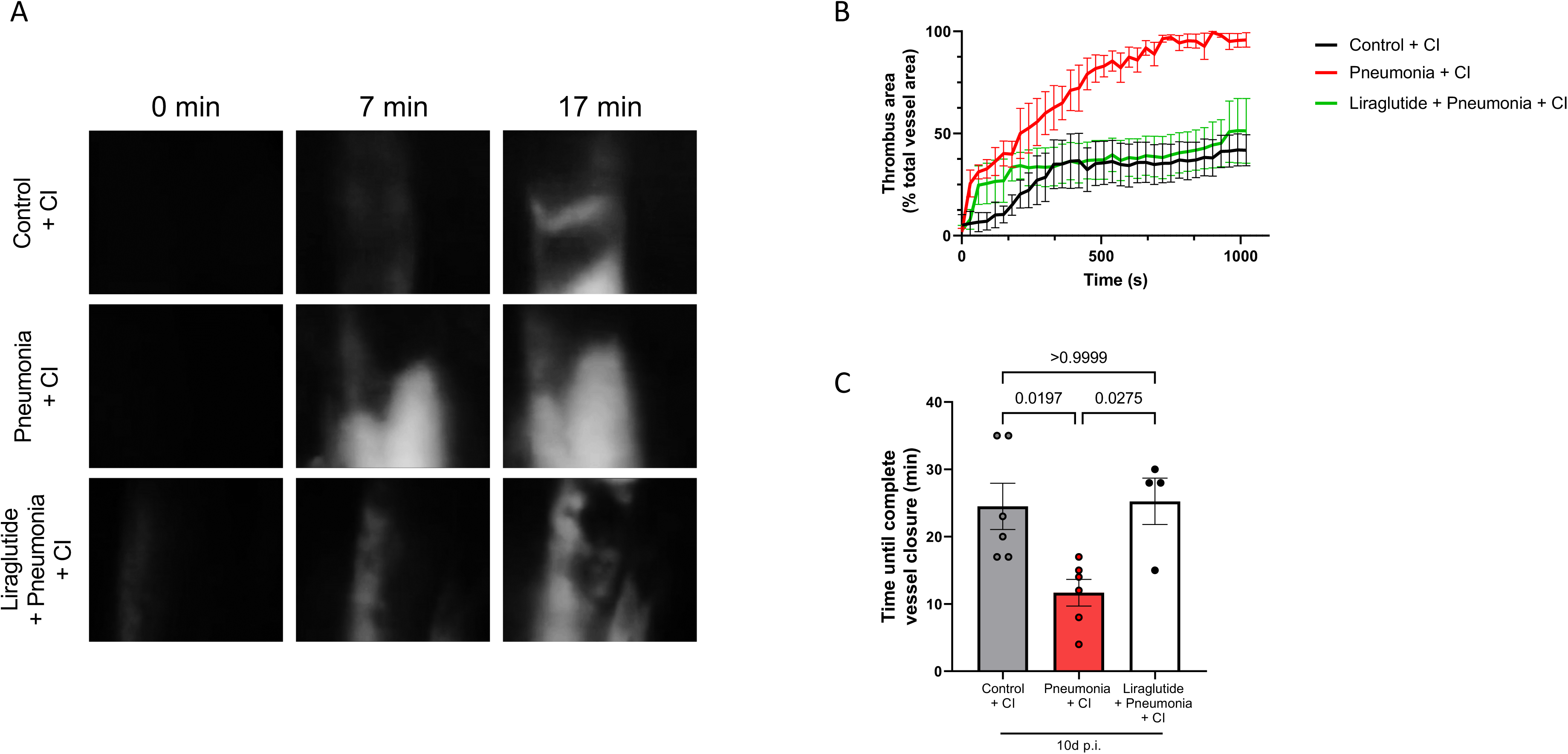
Murine carotid artery intravital microscopy of thrombus formation. **A**, Representative images of thrombus formation in mouse carotid artery induced by FeCl3 in control (n=6), pneumonia (n=6) and liraglutide group (n=4) at three different time points. 10 x magnification. **B**, Time-course of thrombus area. **C**, Plot of time until total carotid artery occlusion.

### Glucagon increases adhesion molecules and monocyte adhesion to the endothelium

Next, we examined whether glucagon promotes endothelial inflammation. To this end, the expression of the adhesion molecules ICAM-1 and E-selectin was measured by flow cytometry. Expression of both adhesion molecules was significantly increased after glucagon stimulation (Figure 7A-C). Consistently, we observed increased adhesion of monocytes to glucagon stimulated HAECs (Figure 7D and 7E). These findings demonstrate a glucagon-induced shift to a pro-inflammatory EC phenotype which likely contributes to impaired endothelial regeneration after injury and increases endothelial thrombogenicity.

**Figure 7.**
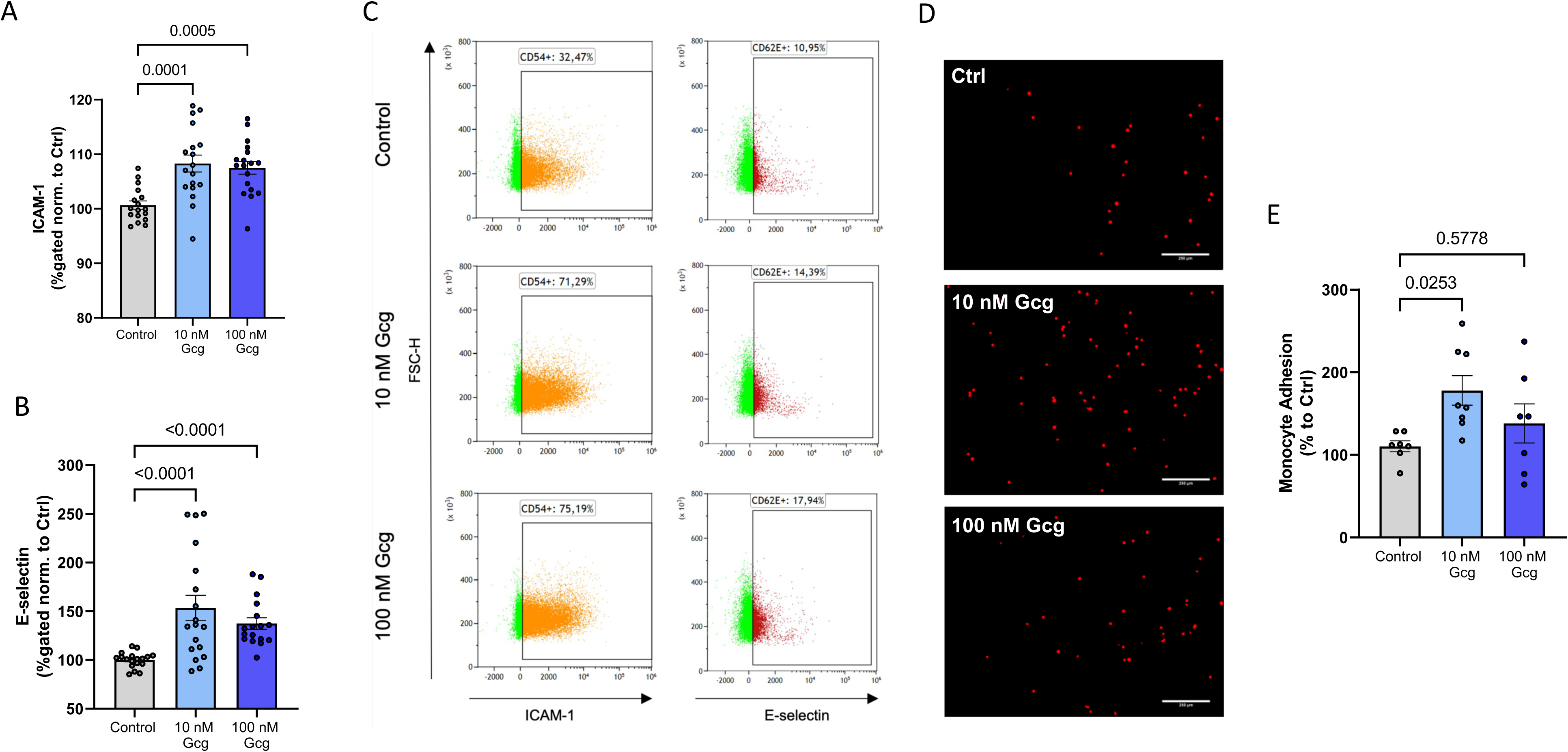
Glucagon increases expression of adhesion molecules and monocyte adhesion on endothelial cells. **A**,**B**, Graphs show changes in expression of ICAM-1 and E-selectin in HAECs stimulated with 10 nM and 100 nM glucagon for 6 hrs compared to control. n=5 per group. The expression of adhesion molecules was measured by flow cytometry. Quantitative data show the percentage of gated CD54^+^ or CD62E^+^ of singlets normalized to control. **C**, Representative flow cytometry plots are shown demonstrating increased levels of ICAM-1 (CD54^+^) and E-selectin (CD62E^+^) expression in HAECs upon glucagon stimulation (middle and lower plot) as compared to control conditions (upper plot). **D**, Representative images of DiI-labeled THP-1 monocytes (red dots) adhered to endothelial cells upon 6 hrs of glucagon stimulation. 10 x magnification, scale bar 250 µm. HAECs subjected to flow conditions with a shear stress of 20 dyn/cm^2^ were stimulated with 10 nM glucagon (n=6), 100 nM glucagon (n=7) or control (n=7) for 6 hrs. **E**, Quantification of THP-1 adhesion to HAECs.

### Lowering plasma glucagon levels reduces endothelial damage

To better understand the potential therapeutic applicability of these findings, we lowered plasma glucagon levels *in vivo* by treating mice with pneumonia additionally with liraglutide, a GLP-1R agonist (0, 24 and 48 hours p.i.) (Figure 8A). The additional administration of liraglutide to infected mice did not alter the lung injury score (Figure S3A and S3B). Notably, mice with additional liraglutide treatment had significantly lower plasma glucagon levels 48 hours p.i., as compared to mice without liraglutide treatment (Figure 8B). Importantly, treatment with liraglutide led to significantly improved vascular regeneration (Figure 8C), suggesting that targeting GLP-1R to lower circulatory glucagon levels, may potentially serve as a promising therapeutic approach to reduce atherothrombotic risk after pneumonia.

**Figure 8.**
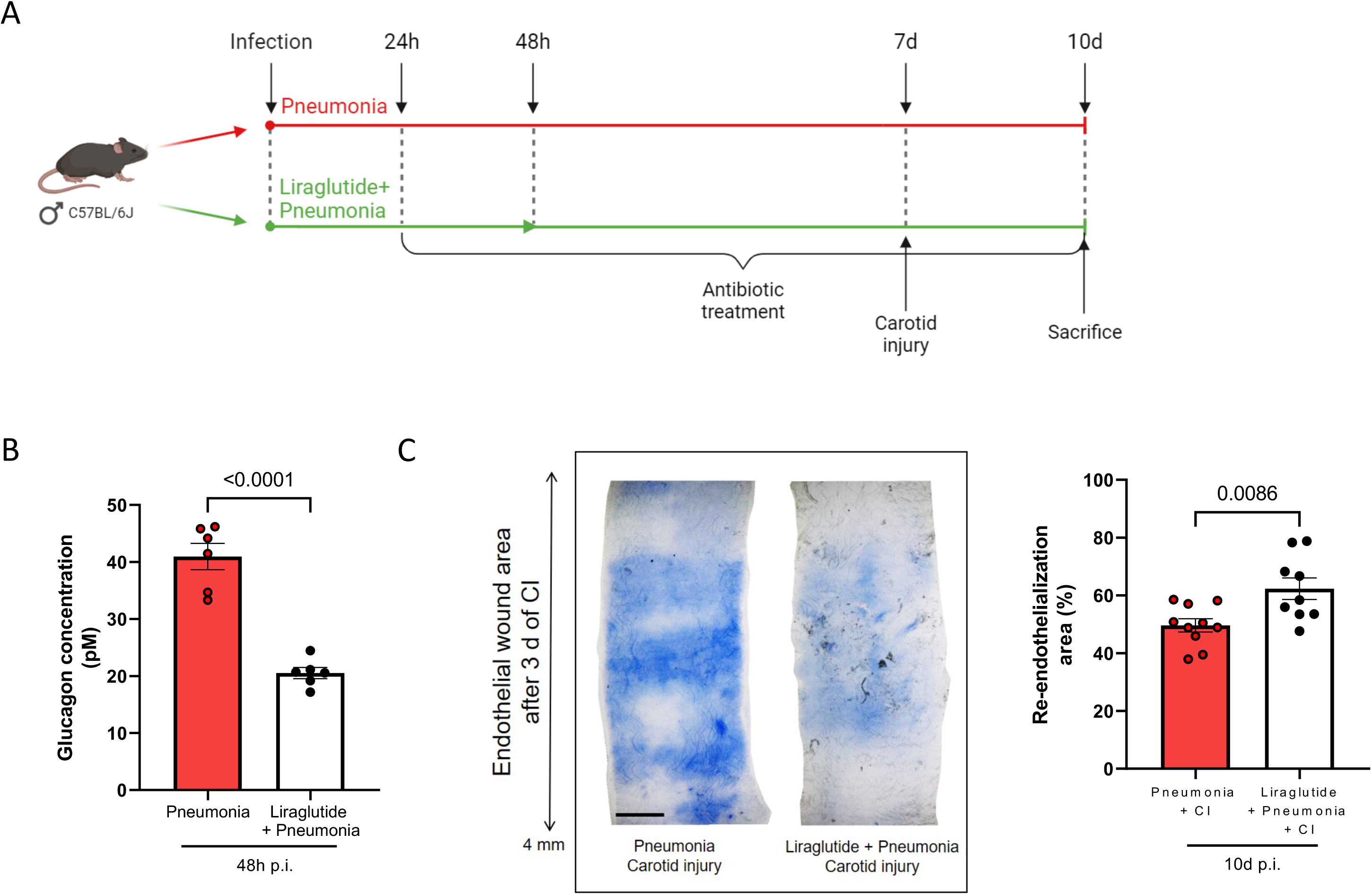
Treatment with liraglutide improves vascular regeneration after injury. **A**, Graphic illustration of experimental setup with liraglutide treatment. Liraglutide (100μg/kg BW) was administered 0h, 24h and 48h p.i. after infection with *S. pneumoniae.* **B**, Blood glucagon concentration measured by ELISA was significantly reduced 48h p.i. in mice treated with liraglutide. n=6 per group. The results of the pneumonia group were the same results used in Figure 2E. **C**, Representative images of Evans blue-stained murine carotid arteries from pneumonia and liraglutide group three days after CI. The blue-stained area corresponds to injured area. 5 x magnification, scale bar=500 μm. n=10 pneumonia. n=9 liraglutide + pneumonia. The results of the pneumonia group were the same results used in Figure 1C.

### Changes of immune cell phenotype in mice with pneumonia

Finally, we examined the immune cell profile in mice 10 days after infection with *S. pneumoniae* when vascular injury was applied by means of flow cytometry. These analyses showed a significant elevation in pro-atherogen Th1 cells (IFNγ^+^/CD4^+^) in the circulatory blood of infected mice (Figure S5A). Interestingly, there were no significant differences in the Th17 (IL17A^+^), Th (CD4^+^) and cytotoxic T cell (CD8^+^) populations in the circulatory blood or in the spleen of mice with pneumonia compared to control mice (Figure S4B-D). Similarly, the frequencies of Ly6C^high^ and Ly6C^low^ monocytes were neither altered in the blood, bone marrow nor spleen of mice with pneumonia (Figure S4E-G). Together these results indicate a substantial resolution of pneumonia-related inflammation at day 10 p.i.

## Discussion

The present study examined the consequences of bacterial pneumonia on endothelial inflammation, thrombogenicity and regeneration after injury. The study provides several important findings: (1) pneumococcal pneumonia impairs vascular regenerative potential after injury; (2) in mice with pneumonia endothelial inflammation and thrombogenicity is increased after vascular injury; and (3) the circulatory level of the peptide hormone glucagon is increased in the resolution phase after pneumonia and may mediate, at least partly, the pathological alterations of the endothelium.

Damage of the vascular endothelium is a characteristic feature in respiratory infections.^15^ While proper endothelial repair mitigates pathologic alterations of the vascular wall, impaired regenerative capacity of the endothelium promotes vascular thrombogenicity^16^ and thus, cardiovascular complications, such as myocardial infarction and ischemic stroke. In the present study, we observed considerably impaired healing capacity of the endothelium in mice with pneumococcal pneumonia. This was accompanied by accelerated thrombus formation at the vascular lesion. Importantly, the levels of monocyte subsets, Th, Th17 and cytotoxic T cells were not different at the time of vascular injury, indicating that the acute inflammatory response had subsided following antibiotic treatment and that the observed vascular effects were related to altered endothelial responses rather than immune cells.

As a potential factor contributing to the impaired healing capability of the endothelium, we identified glucagon to be increased in the circulation of mice. The results of our *in vitro* studies identified detrimental effects of glucagon stimulation on endothelial cell physiology, including impaired migratory capacity and mitochondrial bioenergetic function. Moreover, glucagon treatment promoted inflammatory activation of endothelial cells towards a pro-adhesive phenotype. While previous research on glucagon has focused mainly on the pathophysiology and clinical implications in diabetes^17^ the direct vascular effects of glucagon have been largely ignored so far. However, data from a mendelian randomization study suggest a link between glucagon and ischemic heart disease (IHD) independent from its putative diabetogenic effects.^18^ In this study genetic associations with IHD and type 2 diabetes were obtained for a meta-analysis comprising data from the UK Biobank and the CARDIoGramsplus C4D cohorts and the results demonstrated that genetically predicted glucagon was positively associated with IHD, but not with type II diabetes, nor with fasting insulin, fasting glucose, or glycated hemoglobin. Although, these findings point to a causal role of glucagon in IHD, the detailed pathways through which it affects IHD are yet to be elucidated. Using experimental mouse models and *in vitro* analyses, our current study supports a causal role for glucagon in ischemic events in the context of bacterial pneumonia and provides distinct functional and molecular mechanisms linking glucagon to increased vascular thrombogenicity.

Interestingly, the peak level of glucagon was observed when the mice had recovered from the acute infection, suggesting that the rise in blood glucagon levels does not reflect the increased metabolic demand during the acute phase of infection, but rather occurs as a delayed response. Notably, early suppression of glucagon release by the GLP-1R agonist liraglutide restored the healing potential of injured endothelium and prevented the increased thrombogenicity in mice with pneumococcal pneumonia. However, apart from its glucagon-lowering effect, liraglutide may also exert direct protective effects on endothelial cells. For example, in mice with experimental arterial hypertension, liraglutide was shown to prevent vascular oxidative stress, reduce endothelial NO-synthase uncoupling, and increase NO bioavailability and consequently prevent vascular inflammation.^19^ Moreover, anti-adhesive effects on ECs have been reported for GLP-1R agonists, e.g. through the regulation of Kruppel-like factor 2 (KLF2), a transcription factor known to modulate the inflammatory response of ECs during the development of atherosclerosis.^20^ The limited specificity of liraglutide for glucagon-mediated effects does not take away from the exciting perspective of the potential usefulness of this drug for the prevention of cardiovascular events following pneumonia, which will have to be explored in clinical trials and by analysis of real-life data from large databases in the future.

In conclusion, our observations suggest a novel mechanism linking elevated blood glucagon levels after pneumococcal pneumonia to impaired endothelial regenerative capacity and increased thrombogenicity. Modulation of glucagon release, e.g. by GLP-1R agonists, may provide a novel approach to counteract pneumonia-associated increased atherothrombotic cardiovascular risk. Further clinical studies are needed to evaluate the clinical benefit of targeting glucagon blood level in patients with pneumonia.

## Acknowledgments

The authors would like to thank Ulrike Behrendt, Denise Barthel and Minoo Moobed for excellent assistance.

## Sources of Funding

This study was funded by SYMPATH (“Systems-medicine of pneumonia-aggravated atherosclerosis”, grant number 01ZX1906B), the German Federal Ministry of Education and Research (BMBF), and by grants from the German Heart Research Foundation (DSHF F/01/22) to A.H.

## Nonstandard Abbreviations and Acronyms

ADP: adenosine diphosphate
BW: body weight
CAP: community acquired pneumonia
CI: carotid injury
CVD: cardiovascular disease
DAPI: 4′,6-diamidino-2-phenylindole
DTT: dithiothreitol
EC: endothelial cell
ELISA: Enzyme-linked Immunosorbent Assay
FCCP: carbonylcyanid-4-(trifluormethoxy)phenylhydrazon
FCS: fetal calf serum
Gcg: glucagon
GLP1-R: glucagon-like peptide-1 receptor
HAEC: human aortic endothelial cell
HE: hematoxylin and eosin
ICAM-1: intercellular adhesion molecule 1
IHD: ischemic heart disease
i.p.: intraperitoneal
KEGG: Kyoto Encyclopedia of Genes and Genomes
KLF2: Kruppel-like factor 2
LCA: left carotid artery
NPX: normalized protein expression
OCR: oxygen consumption rates
p.i.: *post infectionem*
PBS: phosphate buffered saline
PFA: paraformaldehyde
*S. pneumoniae*: *Streptococcus pneumoniae*
VCAM-1: vascular cell adhesion molecule 1

## Supplement Figure Legend

**Figure S1.**
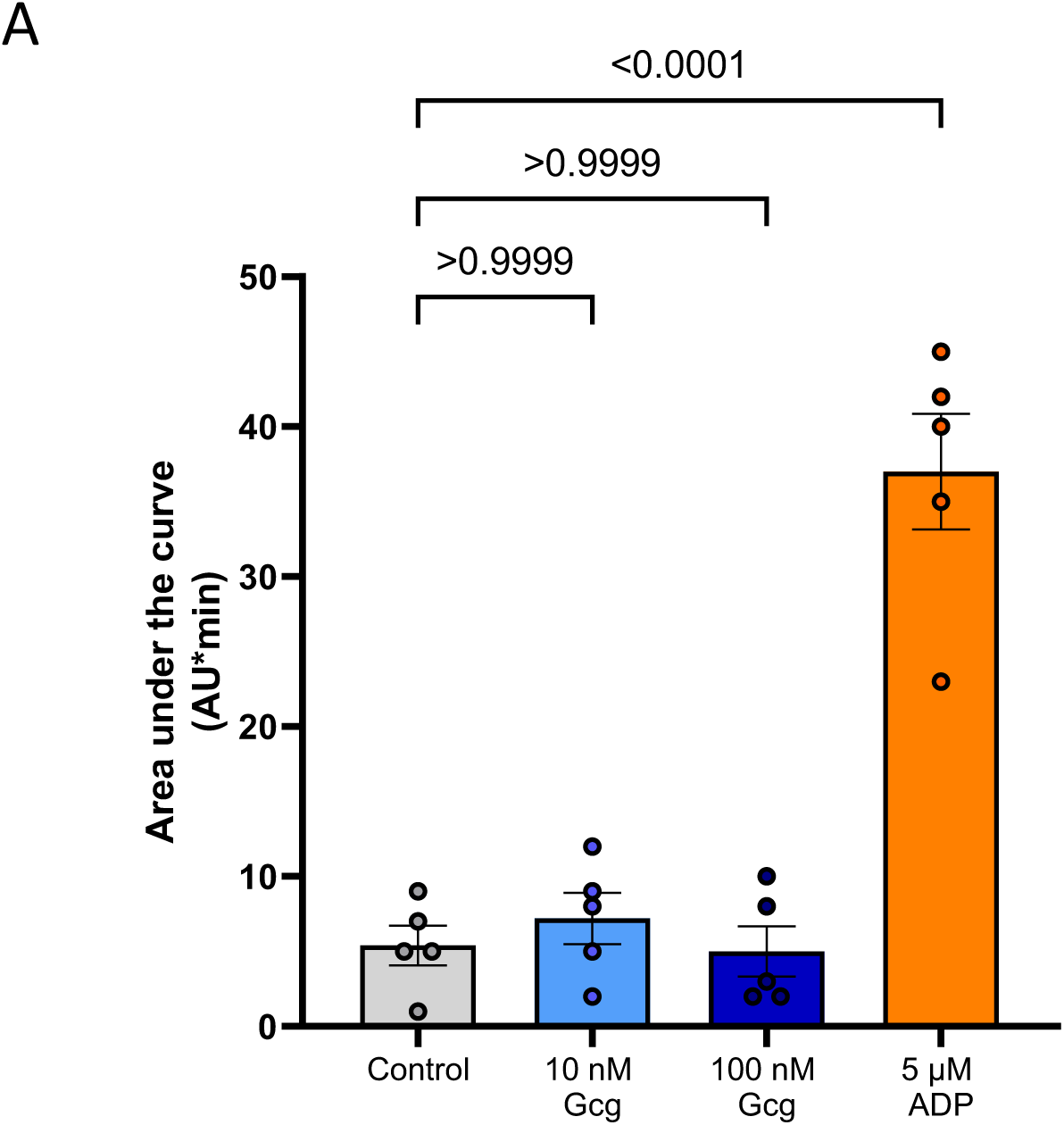
Platelet aggregation in whole blood is not altered by glucagon. **A,** Graph shows platelet aggregation analyses in AU*min per condition. Whole blood samples were stimulated with either 10 nM or 100 nM glucagon, or 1 µm ADP and compared to the control vehicle. n=6 per group

**Figure S2.**
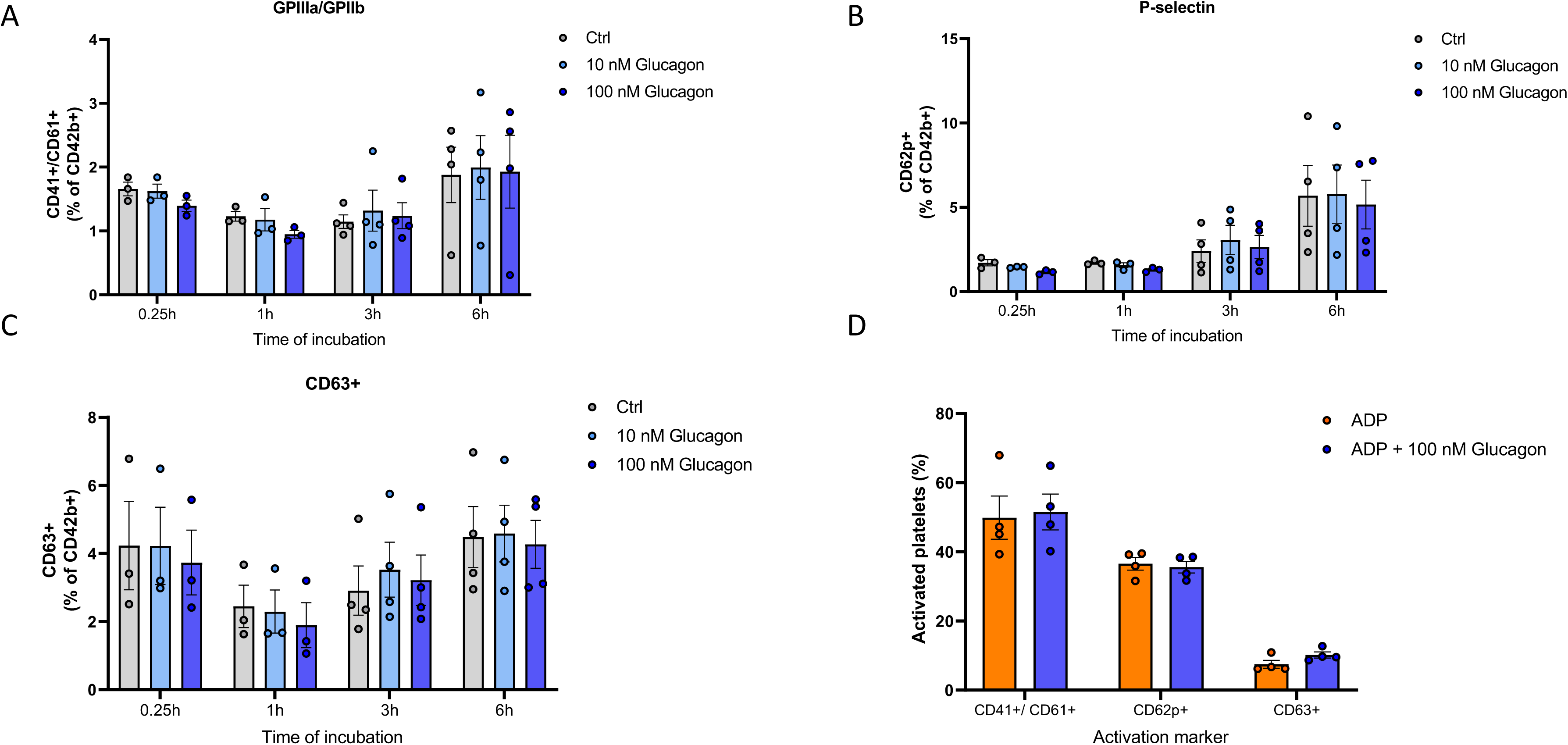
Glucagon does not affect platelet activation markers. **A-C**, The expression of the following markers was examined to determine the activation of platelets in whole blood samples after 10 nM and 100 nM glucagon stimulation for 0.25h, 1h, 3h and 6h compared to control: CD41, CD61, CD62p (P-selectin) and CD63. Quantitative data show the percentage of gated CD41^+^/ CD61^+^, CD62p^+^ and CD63^+^ cells of CD42-positive thrombocytes. **D**, ADP-stimulated expression of platelet activation markers, both with and without 100 nM glucagon stimulation after 6 hrs.

**Figure S3.**
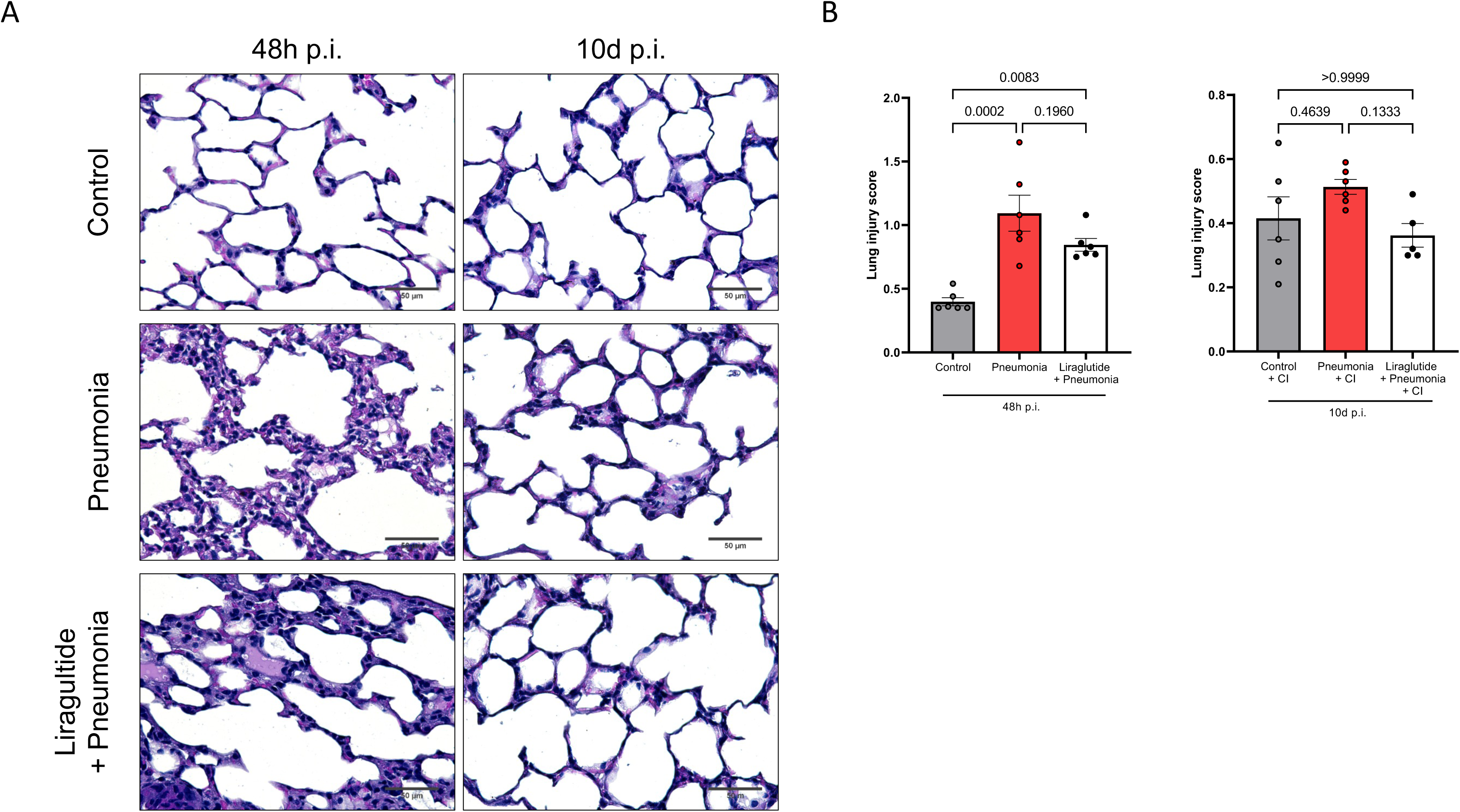
Liraglutide treatment has no effect on lung injury score. **A-B**, Representative images of HE stained lungs from control mice, infected mice without and with liraglutide treatment 48h and 10d p.i. **B**, Quantification of the lung injury score per timepoint. n=4-6 mice per group. 40x magnification, scale bar 50µm. The results of the control mice and pneumonia mice were the same results used in Figure 1B.

**Figure S4.**
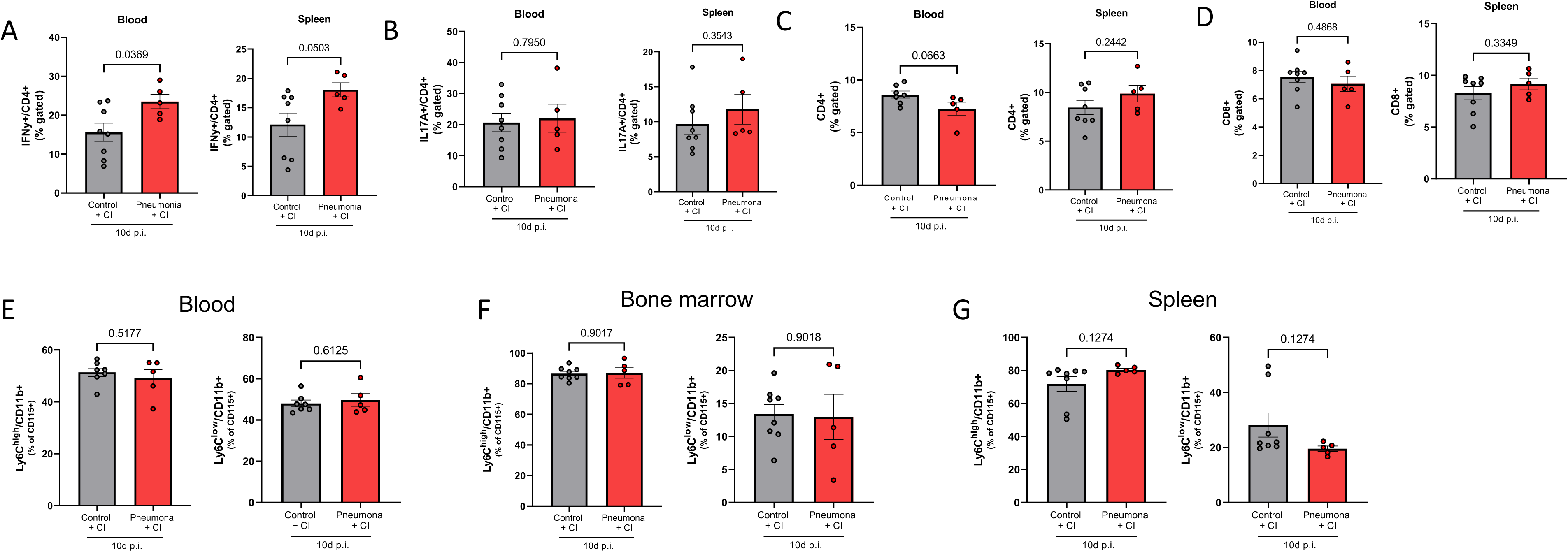
Pro-atherogenic Th1 cells are elevated in mice upon bacterial pneumonia. Flow cytometric analysis of immune cells isolated from murine blood, spleen and bone marrow 10d p.i. **A,** Pro-atherogen Th1 cells are increased in the blood of infected mice as compared to control mice. A trend towards elevation in Th1 cells is shown in the spleen of infected mice. Quantitative data show the percentage of gated Th1 (IFNγ^+^) of CD4^+^ cells. n=8 control. n=5 pneumonia. **B-D,** No alterations were observed in the frequencies of Th17, Th, and cytotoxic T cells in response to pneumonia. Quantitative data show the percentage of gated Th17 (IL17A^+^) among CD4^+^ cells, Th cells (CD4^+^) and cytotoxic T cells (CD8^+^). n=8 control. n=5 pneumonia. **E-G**, Quantitative data show no differences in the percentage of gated Ly6C^high^ and Ly6C^low^ monocytes among CD115^+^ cells in the blood, bone marrow, and spleen between control mice (n=8) and mice with pneumonia (n=5).

## Notes

### Competing Interest Statement

The authors have declared no competing interest.

